# Elucidating the molecular determinants for binding modes of a third-generation HIV-1 integrase strand transfer inhibitor: Importance of side chain and solvent reorganization

**DOI:** 10.1101/2023.11.29.569269

**Authors:** Qinfang Sun, Avik Biswas, Dmitry Lyumkis, Ronald Levy, Nanjie Deng

## Abstract

The first and second-generation clinically used HIV-1 integrase (IN) strand transfer inhibitors (INSTIs) are key components of antiretroviral therapy (ART), which work by blocking the integration step in the HIV-1 replication cycle that is catalyzed by a nucleoprotein assembly called an intasome. However, resistance to even the latest clinically used INSTIs is beginning to emerge. Developmental third-generation INSTIs, based on naphthyridine scaffold, are promising candidates to combat drug-resistant viral variants. Among these novel INSTIs, compound 4f exhibits two distinct conformations when binding to intasomes from HIV-1 and the closely related prototype foamy virus (PFV), despite the high structural similarity of their INSTI binding pockets. The molecular mechanism and the key active site residues responsible for these differing binding modes in closely related intasomes remain elusive. To unravel the molecular determinants governing the two distinct binding modes, we employ a novel molecular dynamics-based free energy approach that utilizes alchemical pathways to overcome the sampling challenges associated with transitioning between two ligand conformations within crowded environments along physical pathways. The calculated conformational free energies successfully recapitulate the experimentally observed binding mode preferences in the two viral intasomes. Analysis of the simulated structures suggests that the observed binding mode preferences are caused by amino acid residue differences in both the front and the central catalytic sub-pocket of the INSTI binding site in HIV-1 and PFV. Additional free energy calculations on mutants of HIV-1 and PFV revealed that while both sub-pockets contribute to the binding mode selection, the central sub-pocket plays a more important role. These results highlight the importance of both side chain and solvent reorganization, as well as the conformational entropy in determining the ligand binding mode and will help inform the development of more effective INSTIs for combatting drug-resistant viral variants.

## 1. Introduction

The HIV-1 strand transfer inhibitors (INSTIs) are important components of antiretroviral therapies (ART)^1–2^. INSTIs work by blocking the catalytic function of the enzyme IN, which assembles into an oligomeric complex with viral DNA known as the intasome and is responsible for inserting the viral DNA into the host cell’s DNA during the early stages of HIV-1 replication. This process is referred to as integration. By inhibiting catalytic integration, INSTIs effectively prevent the establishment of a provirus and further replication within the cell. However, HIV-1 resistance to even the latest clinically used INSTIs is beginning to emerge, as evidenced by accumulating mutations in the IN gene ^3^. In recent years, a novel group of naphthyridine-based third-generation INSTIs have emerged, which demonstrate improved potency against drug-resistant variants of HIV-1^4–5^.

The development of clinically used and developmental INSTIs has, in part, relied upon structural insights gained from intasome assemblies, which are targeted directly by this group of drugs. The structure of the prototype foamy virus (PFV) intasome with bound INSTIs was determined in 2010 ^6^, and this complex has been accordingly used extensively for structure-based drug design efforts ^4, 7–9^. However, recent structures of HIV-1 intasomes with bound INSTIs have revealed that some INSTIs can bind distinctly to intasomes from HIV-1 in comparison to intasomes from the related PFV. Specifically, one of the naphthyridine-containing compounds, 4f (Fig. 1A), which has exhibited superior potency at inhibiting drug-resistant viral variants in comparison to clinically used INSTIs ^4, 10^, stands out by displaying distinct binding modes for these two intasomes (Fig. 1B)^4, 11^. Despite the high degree of similarity in both shape and amino acid composition within the INSTI binding pockets of the two intasomes, the 6-substituted sulfonylphenyl moiety, a critical component that contributes to the potency of naphthyridine compound 4f ^4, 10^, engages in intramolecular stacking with its own naphthyridine core within the PFV binding site of the PFV intasome, leading to a “bent” conformation of the ligand. In contrast, when this same ligand is bound within the pocket of the HIV-1 intasome, the sulfonylphenyl moiety adopts an extended conformation (Fig. 1C). The differences in binding mode lead to distinct interpretations when analyzing mechanisms of drug resistance or when using the structural insights for the purposes of ligand modification, depending on whether the PFV or the HIV-1 model is being used to guide the modeling and ligand optimization.

**Figure 1.**
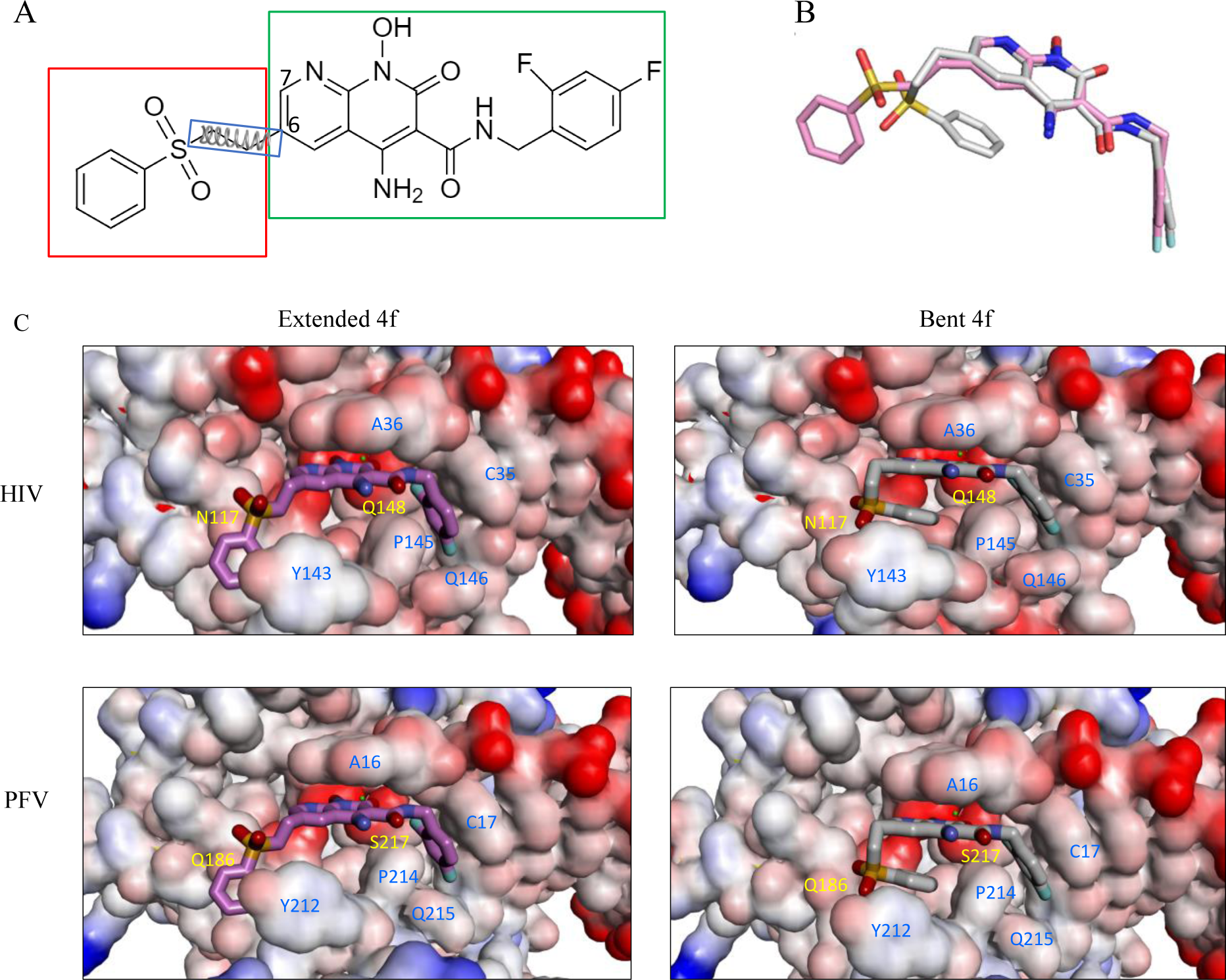
A. Chemical structure of 4f, consisting of a sulfonylphenyl group (red box), a naphthyridine core and a 1,3 difluorobenzene group (green box). The three harmonic dihedral angle restraints that govern the conformation of the sulfonylphenyl group in the R-FEP-R simulations are also indicated. (B) Overlay of two binding modes of 4f: extended (pink) and bent (gray) binding modes. (C). 4f bound to HIV-1 (upper) and to PFV (lower) in the extended (left) and bent (right) conformations.

We were interested in understanding the underlying reasons for the distinct binding modes observed experimentally. Little is known about the physical factors that stabilize the same ligand molecule differently inside binding pockets that are very similar; the RMSD between C-alpha atoms of corresponding binding site residues in the HIV-1 and PFV intasomes are ∼0.5 Å (Table S1, Supporting Information). While variations in the amino acid residues that surround the bound INSTI and DNA substrates are believed to affect the binding of the ligand in the active site, it is unclear which of these residues are most responsible for altering the binding modes. Furthermore, while the structure of the HIV-1 intasome with 4f bound was derived using cryo-electron microscopy (cryo-EM), the structure of the PFV intasome with 4f bound was derived using X-ray crystallography. Whether the observed ligand binding mode is influenced by the conditions under which the experiment was performed (e.g., crystal packing or differences in buffers employed) remains to be examined and helped to motivate this modeling project.

A molecular understanding of the determinants for the INSTI’s binding modes in the two similar viral intasomes could provide insights to help improve inhibitor design. However, accurately estimating the relative thermodynamic stability of different binding modes for a flexible ligand inside a highly packed semi-enclosed space of the binding pocket is known to be computationally challenging^12–15^. While enhanced sampling methods such as metadynamics can be used to connect the two end-state binding conformations via physical pathways^13^, identifying the relevant reaction coordinate to be used in such methods can be nontrivial, as it will involve correlated motions of both ligand and protein side chains to facilitate the conformational transition between the two binding modes inside the binding pocket. The resulting high dimensionality of the conformational free energy surface that needs to be explored can sometimes preclude convergence of the free energy estimation. In contrast to enhanced sampling methods that use pre-defined physical reaction coordinates to connect two conformational basins, the R-FEP-R method (Restrain-Free Energy Perturbation-Release)^16^ uses a dual topology alchemical pathway to circumvent the high free energy barriers that may arise from both intramolecular and intermolecular interactions in a crowded environment. R-FEP-R has been successfully applied to compute the free energy of conformational changes in protein loops^16^ and in DNA base pairing^17^. Here we apply this method to estimate the conformational free energy difference between the two binding modes of the INSTI 4f in complexes with either HIV-1 or PFV intasomes.

The results of the free energy calculations from the R-FEP-R method confirmed that the two distinct conformational binding modes of 4f, one to PFV and the other to HIV, observed using the different experimental techniques, namely cryo-EM and X-ray crystallography, are indeed thermodynamically favored in solution and are not the result of the experimental conditions employed. Analysis of the MD structures revealed that changes induced by 4f binding, in regard to the side chain conformations and the hydration patterns within the central sub-pocket, as well as changes in the torsional entropy in the front sub-pocket of the INSTI binding site, largely explain the relative thermodynamic stabilities of the different binding modes. Based on these structural insights, we performed further R-FEP-R free energy calculations on the different mutants of the two intasomes. The results show that these mutations will indeed significantly change the relative stabilities of the two binding modes in directions consistent with our physical explanation. Our study identifies the important residues responsible for the distinct way that 4f binds to PFV compared with HIV and explains how ligand-induced side chain and solvent reorganization, as well as conformational entropy, affect the mechanisms of binding mode selectivity in the two closely related intasomes.

## 2. Results

### The calculated conformational free energies recapitulate the preference for the experimental binding modes

Previous high-resolution structural biology studies indicated that compound 4f can adopt either a bent or an extended conformation when bound to the active sites of retroviral intasomes^4,11^. As shown in Fig. 1C, the differences in the binding modes of 4f when bound to either the HIV-1 or the PFV intasome are largely confined to the orientation of the sulfonylphenyl group, which is appended to the 6-position of the naphthyridine core, while the remaining parts of the ligand largely superimpose in the two binding modes. The sulfonylphenyl group is connected to the naphthyridine core by three flexible dihedral angles: see Fig. 1A. When the compound 4f is bound to the HIV-1 intasome, the sulfonylphenyl moiety adopts an extended conformation, but when this same compound is bound to the PFV intasome, the sulfonylphenyl moiety adopts a bent conformation and forms an intramolecular stacking interaction with the planar naphthyridine core (Fig. 1C). The shapes of the 4f binding pockets in the two intasomes are very similar (Table. S1, Supporting Information).

In principle, the conformational preferences of the ligand in the two intasome active site pockets may be captured by running long MD simulations initiated from either of the binding modes and if the simulation time is sufficiently long, both binding modes will be sampled according to the Boltzmann factor of their relative free energy. However, this approach of running brute force MD simulations is ineffective because of the high free energy barriers from both intra-molecular and inter-molecular interactions. For example, no binding mode conversion is observed in multiple 50 ns MD simulations starting from either of the binding modes. A much better strategy in these situations is to use enhanced sampling methods like adaptive umbrella sampling^18^ and metadynamics^19^ to overcome the free energy barriers in conformational space. In these approaches, the key is to choose a suitable set of collective variables that defines a transition pathway connecting the two end states; in the present system, it will be particularly difficult for such calculations to converge, because the physical conformational pathways would involve large movements of not only ligand dihedral angles but protein side chain and backbone torsions. As seen in Fig. S1, Supporting Information, to facilitate the conformational transition between the two binding conformations, both the Pro142 backbone and the Y143 side chain must rearrange to avoid steric clash with the sulfonylphenyl moiety of 4f in order to convert from the extended to the bent conformation, and vice versa.

Therefore, to avoid such possible complications, we chose to apply the recently developed R-FEP-R thermodynamic cycle, where the two end states, i.e., the extended and bent conformations of 4f, are connected via an alchemical pathway (Fig. 2).^16^ In this approach, the intermediate system consists of both the extended and bent sulfonylphenyl groups of the ligand 4f using a dual topology, with their relative contributions to the Hamiltonian varied by the continuously varying alchemical λ parameter. In order to observe the alchemical disappearance/reappearance of the two sulfonylphenyl groups, the convergence of the free energy calculation must be facilitated by a symmetrical restrain and release cycle, in which the three dihedral angles in the flexible linker between the sulfonylphenyl group and the 4f core (Fig. 1A) are harmonically restrained to the values corresponding to each of the two end states.

**Figure 2.**
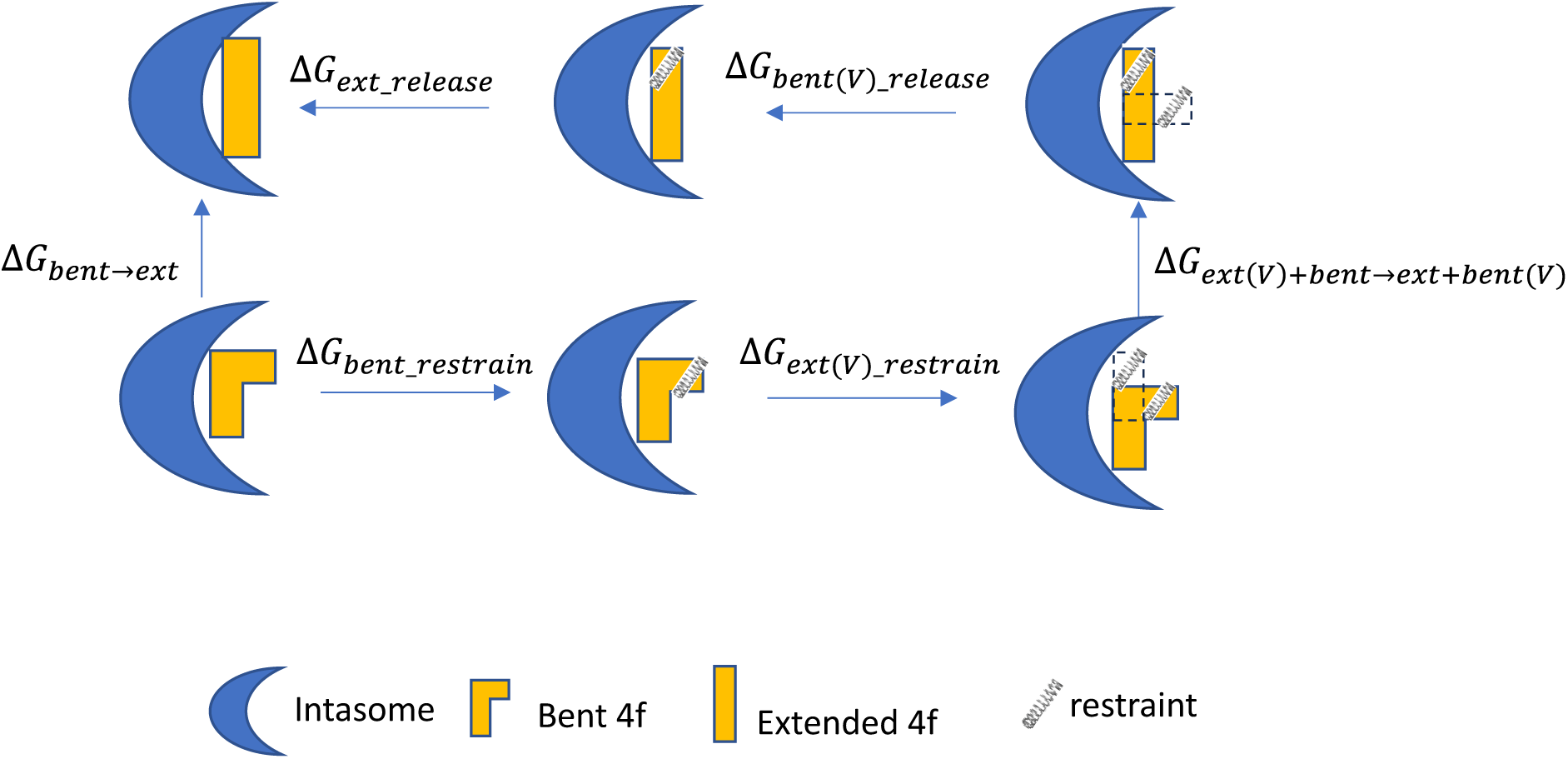
The R-FEP-R thermodynamic cycle for computing the conformational free energy difference of two binding modes. The letter V in the parenthesis indicates the set of atoms in the dual topology set is virtual. The conformational restraints are represented by the springs.

Table 1 shows the R-FEP-R calculated conformational free energy difference 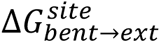. For the wild type (WT) HIV-1 intasome, the extended binding mode is favored with a -2.9 kcal/mol conformational free energy difference relative to the bent conformer. By contrast, when bound to the PFV intasome, the extended 4f conformer is disfavored by 2.2 kcal/mol relative to the bent 4f. Thus, these calculations correctly account for the experimentally observed binding modes, which provides validation for the physical models employed here. The result also supports the inference that the differing binding modes of 4f to intasomes from HIV-1 or PFV are due to differences in the two nucleoprotein complexes, specifically, rather than to the experimental conditions under which the structures were determined, i.e., X-ray crystallography vs. cryo-EM.

**Table 1.** The conformational free energy difference, 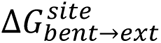, between the bent and extended conformations of 4f in the two intasomes.

| Receptor | | Calculated $\Delta G_{bent \rightarrow ext}^{site}$ (kcal/mol) | Predicted binding mode |
| --- | --- | --- | --- |
| HIV-1 Intasome | Wild Type | $-2.9 \pm 0.4$ | Extended |
| | N117Q | $-2.5 \pm 0.4$ | Extended |
| | Q148S | $-0.7 \pm 0.3$ | Mixed |
| PFV Intasome | Wild Type | $2.2 \pm 0.4$ | Bent |
| | Q186N | $1.4 \pm 0.4$ | Bent |
| | S217Q | $-0.4 \pm 0.2$ | Mixed |

### Understanding the mechanism for the binding mode selectivity in the two intasomes: analysis of MD structures and further R-FEP-R free energy calculations

To gain insights into the molecular determinants that impact the selection of the different binding poses by the two intasomes, we analyzed in detail the MD trajectories of the INSTI binding site in the different 4f binding modes.

In HIV-1 and PFV intasomes, the two distinct conformations of 4f are characterized by the manner in which the sulfonylphenyl group of 4f interacts with two sub-pockets, which we refer to as the front pocket (HIV-1: Asn117/Tyr143/Pro142; PFV: Gln186/Tyr212/Pro211) and the central pocket (HIV-1: Pro145/Gln148; PFV: Pro214/Ser217): see Fig. 3. In the extended binding mode, the sulfonylphenyl group of 4f occupies the front pocket, while the central pocket is filled with solvent. By contrast, in the bent binding mode, the central pocket is partially occupied by the sulfonylphenyl group, leaving the front pocket exposed to the solvent (Fig. 3). Comparing the amino acids at equivalent positions in the two intasomes, we find the following differences: HIV-N117 vs PFV-Q186 in the front pocket, and HIV-Q148 vs PFV-S217 in the central pocket. Analysis of the MD trajectories reveals distinct conformations and hydration patterns involving HIV-Q148 vs PFV-S217 in the central pocket (see also Table 2, Table 3, and Fig. 5). Below we will first discuss the consequences of the HIV-Q148 vs PFV-S217 difference in the central pocket with regard to the binding mode selection, followed by discussing the role of the HIV-N117 vs PFV-Q186 in the front pocket.

**Figure 3.**
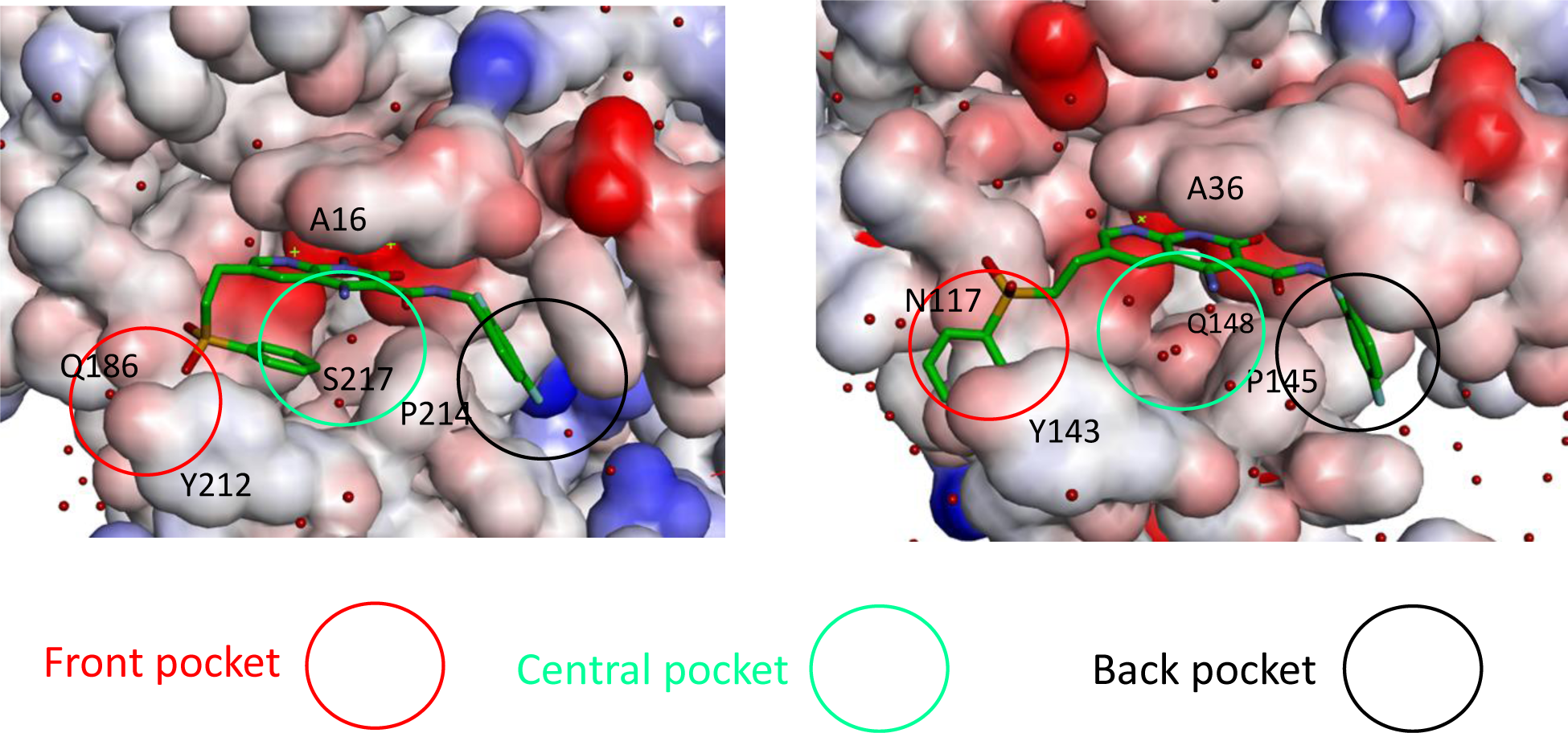
The three sub-pockets (front, central, and back) in the complexes of bent 4f-PFV (left) and extended 4f-HIV-1 (right) INSTI binding sites. The water molecules are shown as red spheres. Note that in the extended binding mode, the sulfonylphenyl group occupies the front pocket, leaving the central pocket solvated, whereas in the bent binding mode, the front pocket is solvated and the central pocket is partially occupied.

**Table 2.** Mean values of the side chain torsion angles of Q148 in the 4f-bound HIV-1 intasome observed in MD simulations, and those in the cryo-EM structure of the apo HIV-1 intasome.

| Structure | $\chi^1$<br>N-CA-CB-CG | $\chi^2$<br>CA-CB-CG-CD | $\chi^3$<br>CB-CG-CD-OE1 |
| --- | --- | --- | --- |
| 4f-bound (extended) | -66.0° | 91.5° | 42.8° |
| 4f-bound (bent) | -59.7° | 145.7° | -120.1° |
| Apo | -49.3° | 103.4° | 29.5° |

**Table 3.** Average number of water molecules occupying the central cavity observed in MD trajectories.

| Intasome | extended 4f | bent 4f |
| --- | --- | --- |
| HIV | 5-6 | 0 |
| PFV | 6-7 | 1-2 |

### The central sub-pocket is primarily responsible for the binding mode selection by the two intasomes

We first consider how the compound 4f engages the HIV-1 intasome in its two distinct binding modes. In the extended binding mode, the sulfonylphenyl group of 4f occupies the front pocket and forms favorable intermolecular π−π stacking interactions with Tyr143 of HIV-1 (Fig. 3). In this binding mode, the central pocket is occupied by solvent, allowing the side chain of Gln148 to adopt a conformation close to the apo-state conformation (Table 2 and Fig. 4) and is well solvated by ∼6 water molecules (Table 3 and Fig. 3). In the bent binding mode, the sulfonylphenyl group folds back and partially occupies the central pocket, making a favorable intramolecular π−π stacking interaction with the naphthyridine ring system of 4f; however, the favorable intermolecular π−π stacking with the front pocket residue Tyr143 is lost. Importantly, the central pocket is now partially occupied by the sulfonylphenyl moiety of 4f, leading to its nearly complete desolvation. As the number of water molecules hydrating the central pocket drops from ∼5-6 to zero, the resulting loss of favorable interactions between water and the polar side chain of Gln148 substantially destabilizes the bent binding mode. Furthermore, the partial occupation of the central pocket by the sulfonylphenyl group of the bent 4f causes the Gln148 side chain to rearrange and adopt a high free energy conformation that is distinct from that of the apo-state (see Table 2, Fig. 4 and Fig. 5A). As we have previously shown^20^, such ligand-induced conformational reorganization in the receptor is generally associated with a free energy penalty and results in decreased thermodynamic stability of the corresponding ligand-bound state.^12, 20–22^

**Figure 4.**
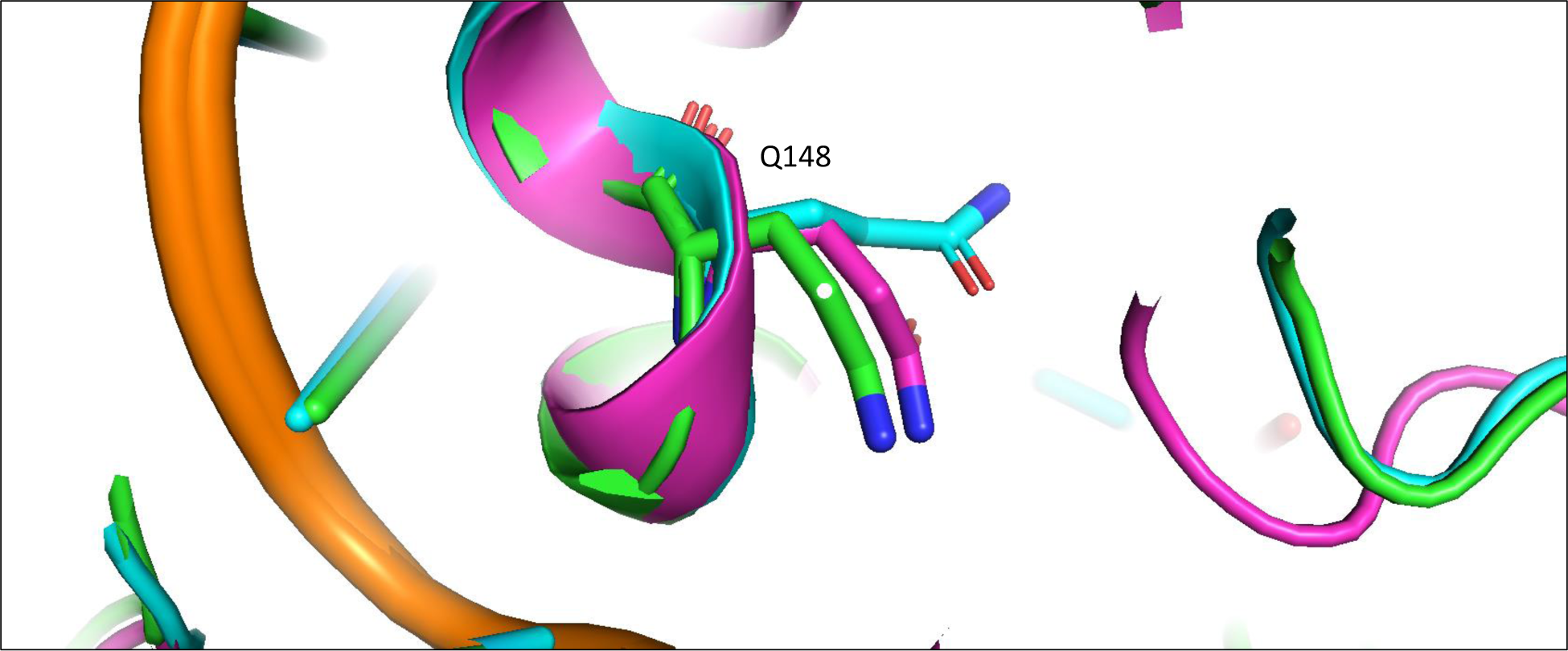
The rotamer states of Q148 of HIV-1 in the apo and 4f bound states. pink: apo; green: extended 4f bound; blue: bent 4f bound.

We next consider how compound 4f engages the PFV intasome in its two distinct binding modes. As in the 4f-HIV-1 complexes discussed above, the extended binding mode is stabilized by the intermolecular π−π stacking between the sulfonylphenyl group and the front pocket residue in this case Tyr212 of PFV, whereas the bent binding mode is primarily stabilized by the favorable intramolecular π−π stacking interaction between the sulfonylphenyl group and the naphthyridine ring system of 4f. However, unlike the 4f-HIV-1 complexes, here the difference in the protein environment within the central pocket in PFV shifts the thermodynamic balance in favor of the bent binding mode. In the PFV intasome, the residue corresponding to HIV-1 Gln148 is Ser217 (Fig. 1C). Serine has a shorter and less polar side chain in comparison to glutamine. As a result, when 4f engages the PFV intasome in the bent conformation, the central pocket residue Ser217 can maintain the same conformation that is observed when 4f engages the PFV intasome in an extended conformation (Fig. 5B). In addition, because of the smaller size of the sidechain of Ser217, the central pocket remains weakly solvated by 1-2 water molecules even when the sulfonylphenyl group of the bent 4f partially occupies this space (Table 3 and Fig. 3). Thus, the bent 4f in the PFV intasome active site is expected to lead to a smaller conformational reorganization free energy, and to a smaller desolvation free energy cost when compared with that in the HIV-1 intasome, where the corresponding central pocket residue is Gln148.

**Figure 5.**
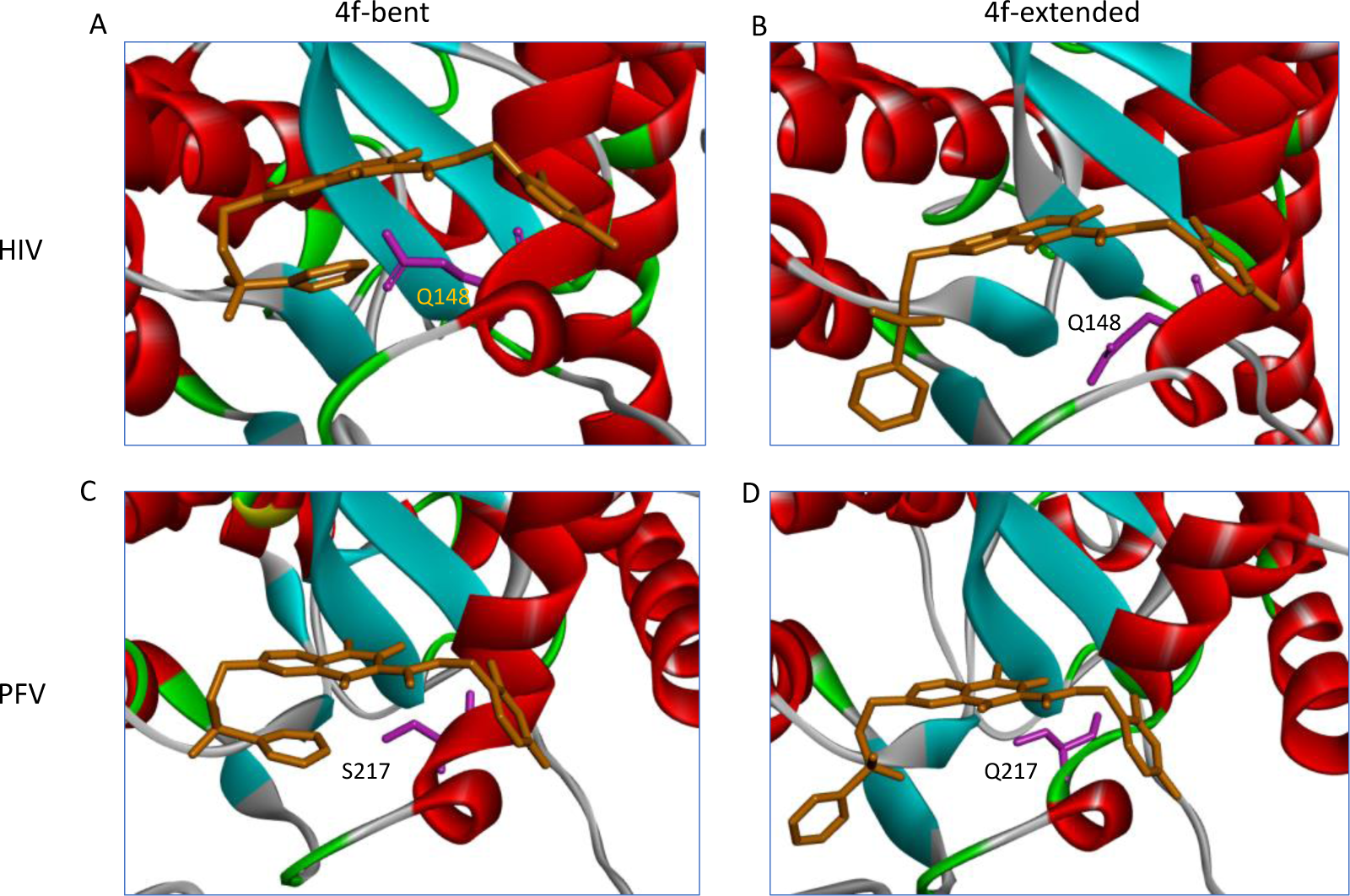
Upper: conformations of the Gln148 in the extended (A) and bent (B) binding modes in the HIV-1 intasome. Lower: conformations of the Ser217 in the extended (C) and bent (D) binding modes in the PFV intasome.

Overall, these analyses suggest that there are two primary determinants governing the binding mode selection by the HIV-1 and PFV intasomes: (i) there is a desolvation free energy cost to the bent conformation of 4f in the HIV-1 intasome when the central pocket residue is Gln148; by contrast, there is a smaller free energy penalty for desolvation in the case of PFV, where the corresponding residue is Ser217. (ii) there is also an associated side-chain reorganization of Gln148 in HIV-1, which is not observed in the corresponding residue Ser217 in PFV. Collectively, these observations explain why the extended binding mode is favored in the 4f-HIV-1 intasome complex, whereas the bent binding mode is more stable in the 4f-PFV intasome complex, as captured via structural biology.

To test our hypothesis that the central pocket determines the binding mode preference, we generated *in silico* substitutions and performed R-FEP-R calculations on the Q148S mutant of the HIV-1 intasome and the S217Q mutant of PFV (Table 1). If Q148 in HIV-1 is indeed causing the extended binding mode to be more stable, then mutating this residue to a Serine, which is the corresponding residue in PFV IN, is expected to significantly reduce the relative thermodynamic stability of the extended binding mode over the bent mode. Similarly, if S217 in PFV is primarily responsible for the binding mode selectivity to favor the bent mode, then mutating this residue to a Glutamine should make the extended binding mode substantially more stable. Table 1 shows the results of R-FEP-R calculations on these mutants. In the HIV Q148S mutant, the calculated free energy difference 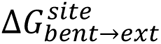 still favors the extended binding mode, but the calculated free energy difference is just - 0.7 ± 0.3 kcal/mol which is significantly smaller than the 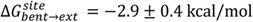 observed with the WT HIV-1 intasomes. In fact, since the value of 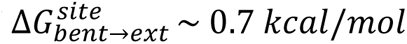 for the HIV Q148S translates to a population ratio of extended and bent 4f of ∼3:1, we accordingly designate these as “mixed” populations. Similarly, in the R-FEP-R simulation of the PFV S217Q mutant, switching the serine to glutamine caused a more dramatic change in free energy, such that the binding mode preference was actually reversed to slightly favor the extended binding mode over the bent binding mode. In both cases, the single point mutation causes significant changes in the binding mode stability by ≥ 2.2 kcal/mol, in the expected directions. These free energy results therefore support the conjecture that the Gln148(HIV-1)/Ser217(PFV) pair in the central pocket is a key factor in determining the binding mode preference in the two intasomes.

### The front sub-pocket makes a comparatively smaller contribution to the binding mode selection by the two intasomes

In addition to identifying the important role of central pocket residues Gln148 (HIV-1) and Ser217 (PFV), we also examined how the front pocket residues Asn117 (HIV-1) and Gln186 (PFV) impact the binding mode preference. In the extended 4f binding mode, the Cβ atoms of Asn117 of HIV-1 and Gln186 of PFV form a similar nonpolar interaction with the sulfonylphenyl ring (see Fig. S2, Supporting Information), whereas in the bent binding mode, such interactions are absent, because the front pocket is unoccupied. Importantly, the length of the side chain matters and Gln186 (PFV) is longer than Asn117 (HIV-1) by one rotatable bond. Consequently, when the ligand 4f binds in the extended binding mode, placing the sulfonylbenzene ring into the front pocket, the side chain of Gln186 of PFV experiences a larger ligand-induced conformational entropy loss than does the side chain of Asn117 of HIV-1. This can be observed by the side chain torsion angle distributions observed in the MD trajectories (Fig. 6 and Fig. 7). Specifically, in the extended binding mode, the conformation of the side chain dihedral angle χ2 of Gln186, PFV, is restricted and resides within a single basin (Fig. 6, upper panel), but in the bent binding mode, it samples all three conformational basins (Fig. 6, lower panel). This translates to a free energy contribution of Δ*G_bent→ext_* (Q186 − PFV) ≈ −*KT* ln 3 = 0.65 kcal/mol, i.e., destabilizing the extended binding mode in PFV relative to the bent binding mode.^23^ In HIV-1 however, the corresponding residue is Asn117, which has a shorter side chain. Asn117 largely occupies a single dominant basin in both extended and bent binding modes of 4f (Fig. 7), giving rise to a smaller entropy loss that does not significantly impact the relative thermodynamic stability of the two 4f binding modes.

**Figure 6.**
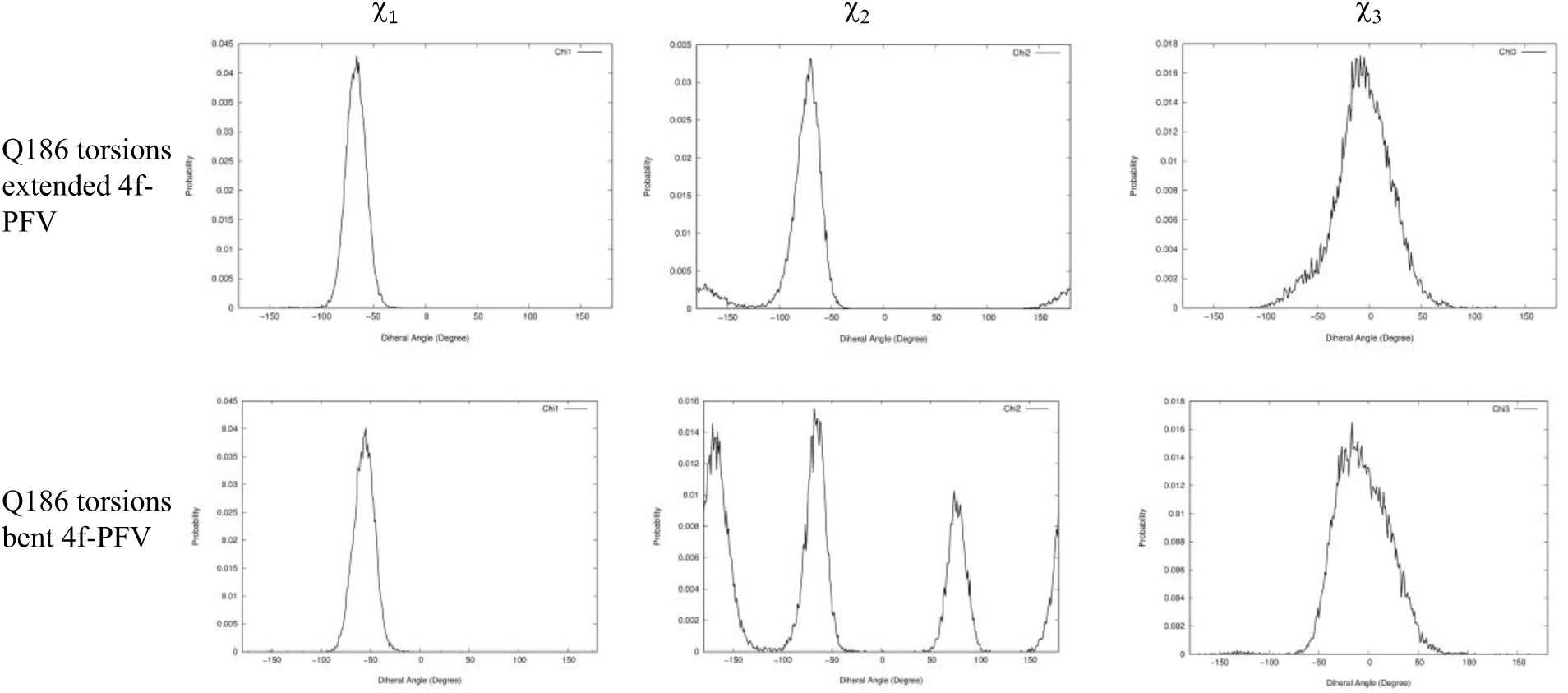
Distributions of side chain dihedral angles of Q186 of PFV in the two 4f binding modes. All data are extracted from 30 ns MD simulations of the corresponding complexes.

**Figure 7.**
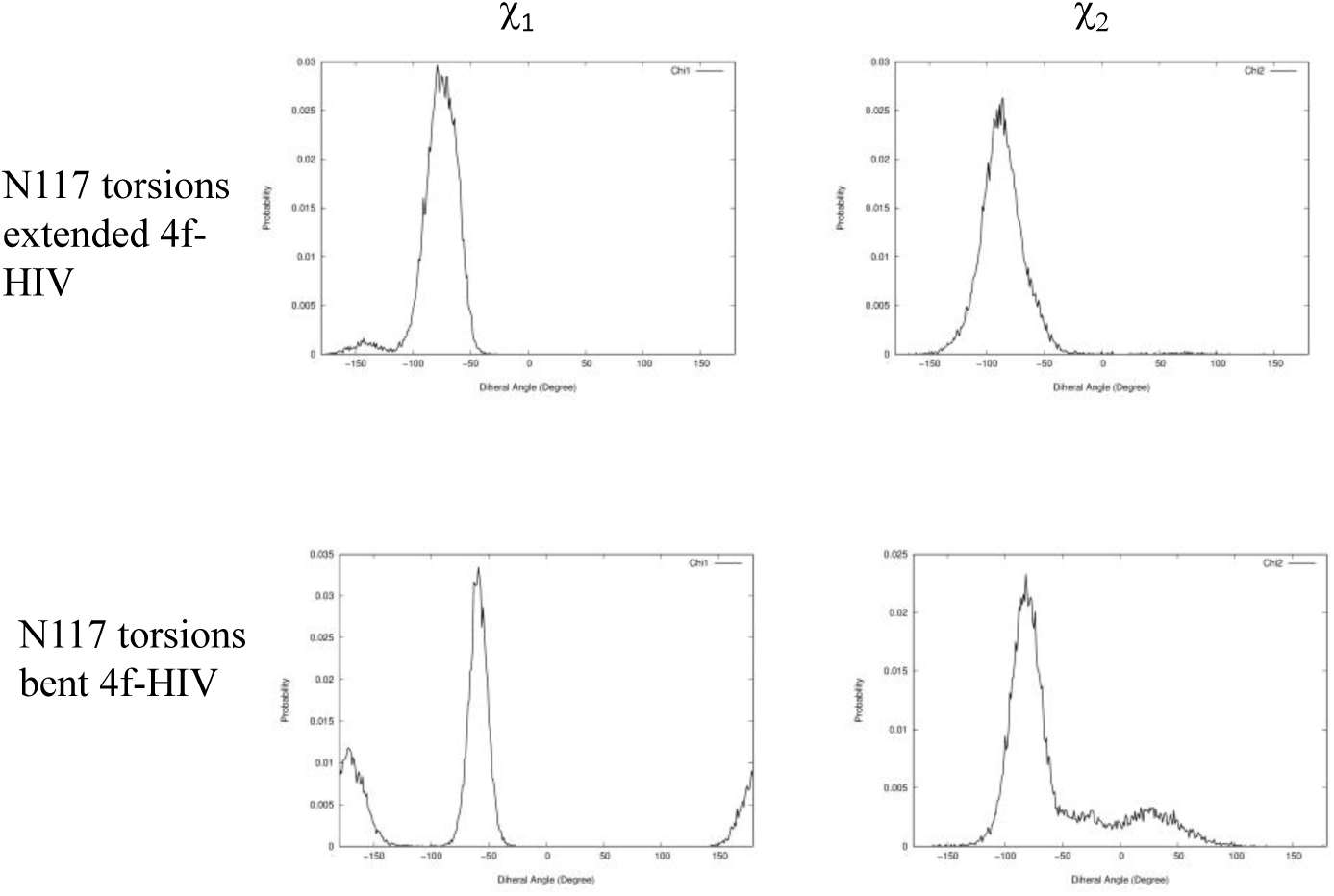
Distributions of side chain dihedral angles of N117 of HIV in the two 4f binding modes. All data are extracted from 30 ns MD simulations of the corresponding complexes.

To further probe the relevance of the residues within the front pocket, we performed R-FEP-R free energy calculations on both the HIV N117Q mutant and the Q186N mutant of PFV (Table 1), both mutations are in the front pocket. In the HIV N117Q mutant, the conformational free energy difference 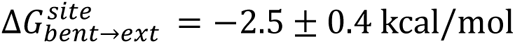, compared to 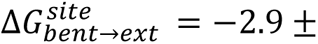 0.4 kcal/mol for the wild-type HIV. Thus, mutating the Asn117 to a Glutamine indeed destabilizes the extended binding mode of 4f by ∼0.4 kcal/mol, as predicted by our analysis based on the torsional entropy analysis. Similarly, in the PFV Q186N mutant, the bent binding mode is destabilized by ∼0.8 kcal/mol by mutating Gln186 to Asn, again consistent with our entropy analysis of Gln186 vs N117. These results, therefore, suggest that, in addition to the prominent role of the central pocket residues Gln148 (HIV-1) and Ser217 (PFV), the distinct residues Asn117 (HIV-1) or Gln186 (PFV) in the front pocket also contribute and help to explain the binding mode selection by the two intasomes, even though this effect is smaller than the effects induced by the mutations (HIV-1 Q148S, and PFV S217Q) in the central pocket.

## 3. Discussion

### Implications of the findings for drug resistance and ligand binding

The findings here have relevance to our understanding of the role of mutations in and around the active site of IN for drug resistance and ligand binding. For example, our simulations suggest that the mutation N117Q in HIV-1 IN, which would be expected to yield viable viruses without significantly compromising enzyme activity^24–25^, may be a DRM candidate in future selection experiments that employ naphthyridine-based compounds or other ligands^26^ containing chemical extensions that protrude into the solvent-exposed cleft of the intasome. N117Q will be more conformationally restricted and thus is expected to lead to an entropic loss when larger compounds, such as 4f, are bound to the intasome. Future selection experiments will test this idea. Furthermore, our observation that the residue at position HIV-1 IN_148_ influences ligand binding to a greater extent than other residues was unexpected and revealing. Q148H/K/R are prominent DRMs, arising frequently in viruses derived from patients on both 1^st^ and 2^nd^ generation INSTI therapy^27–30^. Although the mutation Q148S does not arise in HIV-1 IN (here, it was tested solely for the purpose of comparing with the analogous mutation in PFV IN), the insights gained from our simulations strongly suggest that the nature of the residue at this position will affect ligand binding and selectivity. Specifically, mutations Q148H/K/R should differentially influence the local hydration pattern in pocket 2. Indeed, an analysis of the previously solved PDB structures of the HIV-1 intasomes containing the mutation Q148H (PDB: 8FNP), Q148K (PDB: 8FNN), and Q148R (PDB: 8FNO), indicates that there are distinct hydration patterns within pocket 2. This implies that a careful analysis of the water profile, specifically in pocket 2 but also the network as a whole, will be important for future structure-based drug design, as both the configuration and the total number of waters will affect the free energy of ligand binding.^31–32^ These insights will extend to the development of 3^rd^ generation INSTIs^26, 33–34^.

### Atomic models of HIV intasomes for studying INSTI binding and drug resistance

The first INSTIs were introduced into the clinic in 2007 ^35^. A mechanistic understanding of their mode of inhibition was elucidated in 2010 with the experimental structures of PFV intasomes containing bound INSTIs^6^. Since then, the PFV intasome has been used extensively for structure-based drug design^4, 7–8, 36–37^, including for the development of second-generation clinically used inhibitors, and for rationalizing mechanisms of drug resistance^9^. However, work by two independent laboratories recently revealed that the PFV intasome has limitations and is too divergent from the HIV-1 intasome to properly interpret mechanisms of drug resistance^33^ or to employ in structure-based drug design^34^. On the latter point, we previously observed that the exact same ligand can engage two different intasomes distinctly, but we could not explain the underlying basis of this differential binding mode. Here, we provide such an explanation. Given the increasing emphasis on understanding mechanisms of HIV-1 resistance to therapy and on targeting, specifically, drug resistant viral variants using novel INSTIs^10, 26, 38^, it is thus important to now use experimental structural biology data derived from HIV-1 intasomes for rationalizing mechanisms of resistance and as starting points for structure-based drug design. The growing accumulation of structural biology data defining the conformations of HIV-1 intasomes, including both WT and DRM variants bound to both clinically used and developmental INSTIs, will help to this end.

### Utility of the alchemical R-FEP-R for predicting ligand binding modes in challenging environments

The utility of the alchemical R-FEP-R method for predicting ligand binding modes in challenging environments is demonstrated in this study. We show that this method can accurately compute the relative stabilities of ligand binding modes in the challenging binding site environments found in HIV-1 and PFV intasomes, where ligand-induced side chain and solvent reorganization, as well as configurational entropy, play significant roles.

As observed in Table 1, R-FEP-R calculations are capable of accounting for the relative thermodynamic stabilities of binding modes that differ by approximately 2 kcal/mol, exceeding the accuracy expected from more approximate methods like MM-PB(GB)SA calculations. Compared to physical pathway-based free energy methods such as umbrella sampling or metadynamics, the use of alchemical dual topology in R-FEP-R simplifies the formidable task of choosing reaction coordinates and/or overcoming the large free energy barrier that separates the binding modes. In our work, the two binding modes of 4f only differ in their dihedral angles. However, it’s important to note that different binding modes can involve differences in both internal parameters (such as torsions) and external factors (i.e., ligand translation and orientation). Importantly, the R-FEP-R method is well-suited to address the free energy difference between binding poses that also involve ligand external degrees of freedom. This is achieved by introducing appropriate restraints on ligand translation and orientation in the dual topology setup of the hybrid system. Consequently, we anticipate that the R-FEP-R method will serve as a general solution for rigorously calculating the relative thermodynamic stabilities of multiple ligand binding modes.

## 4. Conclusion

We have investigated the mechanism responsible for the two distinct conformations of a third-generation INSTI 4f in complex with the PFV and HIV-1 intasomes, associated with different binding modes to these viral enzymes, by applying a novel molecular dynamics-based conformational free energy method R-FEP-R. The calculated conformational free energy differences between the two binding modes predict that the extended 4f conformation is favored in the HIV-1 intasome relative to the bent conformer, whereas the bent 4f conformation is favored in the PFV intasome. Both results agree with experimental structural observations, obtained from different sources. Our analysis of the MD simulations of the different binding modes in the two closely related intasomes, supported by additional R-FEP-R free energy calculations which probe the effects of mutations, has identified the crucial role of the central pocket residue Q148 (HIV-1) and S217 (PFV) and, to a lesser extent the front pocket residues N117 (HIV-1) and Q186 (PFV), in determining the binding modes of 4f in the two intasomes. This study highlights the importance of ligand-induced protein side chain and solvent reorganization, and the conformational entropy change in ligand binding and helps inform further development of third-generation INISTIs to combat the rapidly emerging HIV-1 variant strains resistant to existing INSTIs.

## 4. Methods and Materials

### System Setup and Simulation Details

The starting structure of the HIV-1−4f complex is taken from the Cryo-EM structure of compound 4f bound to the HIV-1 intasome (PDB 6PUZ)^34^ and the starting structure PFV−4f is from the crystal structure of 4f bound to PFV intasome (5FRO)^4^.

Molecular dynamics simulations were performed with GROMACS version 2020.3^39^. The HIV-1 and PFV intasome proteins were modeled with the ff19SB force field^40^, the DNA was modeled using the Parmbsc1 force field^41^, and the ligand with the General Amber force field (GAFF)^42^ and the AM1-bcc charge models.^43^ The accuracy of the force field model can have a significant impact on the calculated free energy.^44–46^ Here, to properly model the metal ion Mg^2+^ coordination with the protein and ligand, we use a bonded model involving bond, angle, dihedral, electrostatic, and van der Waals terms^47–48^ that are parameterized by following the procedure developed by Pengfei Li and Merz groups (Metal Center Parameter Builder (MCPB)). The bonded models^49^ treat the interactions between the metal ion and its ligating atoms as covalent bonds. Lin and Wang^50^ applied the bonded model through Seminario method^51^ (using the Cartesian Hessian matrix to calculate the force constant) to zinc complexes with the general AMBER force field (GAFF) and demonstrated that the effectiveness of the bonded model performed in structural optimizations and molecular dynamics simulations of four selected model systems. The benchmarking simulations by Melse and Zacharias^52^ applying the MCPB.py program reproduced the Zn^2+^ sites accurately, including mono- and bimetallic ligand binding sites. To compute force field parameters for the Mg2+ associated atoms of IN (Fig S3), we have applied MCPB.py program with GAMESS-US to generate the bond, angle, corresponding force constants, the partial charge, and the VDW parameters (Table S2).

One intasome unit was placed at the center of a rectangular box and the distance between the solute and the edge of the box was set to be ≥ 1.0 nm. A 9.4 nm × 12.7 nm × 10.2 nm box containing 30,200 OPC water molecules and 4 Na^+^ is used for HIV-1 and an 11.7 nm × 11.1 nm × 14.0 nm box containing 48,800 OPC water molecules and 17 Na^+^ is used for PFV. The total number of atoms in the simulation box is ∼132,000 for HIV-1 and 206,000 for PFV. Additional Na^+^ and Cl^−^ ions were added to obtain 0.15 M NaCl in the simulation box.

The conformational free energy difference between the extended and bent binding modes of 4f is computed using the R-FEP-R method (Fig 2),^16, 53^ which uses an alchemical pathway instead of the traditional physical pathway approach to connect the two conformational basins. In this method, the ligand atoms are divided into the shared part and the varying segment (see Fig. 1A), with only the latter participating in the conformational change. R-FEP-R uses the dual topology approach in which the hybrid system contains atoms from both end-states with their contributions to the Hamiltonian controlled by a varying λ value. The free energy difference is computed as the sum of the following components, i.e.,

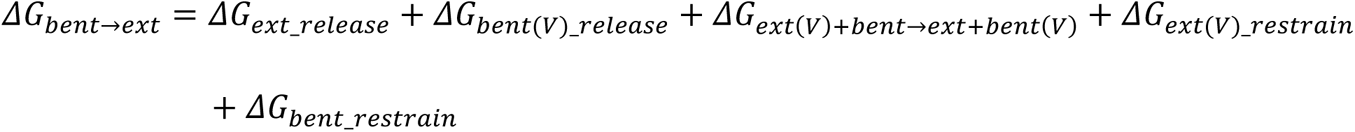

Here Δ*G_bent_restrain_* and Δ*G*_*ext(V)_restrain*_ are the free energies of restraining the real and virtual segments in the initial state (bent), while Δ*G*_*bent(V)_release*_ and Δ*G*_*ext_release*_, are the free energies to release the restrains on the virtual and real segments in the final (extended) state. When the force constants of the restraints used are large, the Δ*G*_*ext(V)_restrain*_ and Δ*G*_*bent(V)_release*_ are equal in magnitude (and opposite in sign) and cancel out with each other.^16, 53^ Δ*G*_*ext+bent(V)→bent+ext(V)*_ is the free energy of alchemically converting the virtual segment in the initial state into the real one in the final state (under conformational restraints). All three free energy components Δ*G*_*ext+bent(V)→bent+ext(V)*_, Δ*G*_*ext_release*_, and Δ*G*_*bent_restrain*_ are computed using FEP.

For calculating Δ*G*_*ext+bent(V)→bent+ext(V)*_, 39 alchemical λ windows were used: λ = 0 0.000001 0.000002 0.000005 0.00001 0.00002 0.00005 0.0001 0.0002 0.0005 0.001 0.002 0.004 0.01 0.03 0.06 0.1 0.15 0.2 0.29 0.38 0.46 0.54 0.62 0.71 0.8 0.85 0.9 0.94 0.97 0.99 0.996 0.999 0.9995 0.9999 0.99995 0.999999 0.999 9999 0.99999999 1.0. At each λ, the equilibration was performed for 1 ns first. The production run was then performed for 10 ns at 300 K. The constant temperature was maintained by the modified Berendsen thermostat^54^. The electrostatic interactions were treated with the particle mesh Ewald (PME)^55^ method with a real-space cutoff of 1.0 nm. MD simulations were performed with a time step of 2 fs and energy files were saved every 2ps.

The number of alchemical λ windows used for computing the release/restraint free energies Δ*G*_*ext_release*_ and Δ*G*_*bent_restrain*_ was set to be 23 (λ = 0 0.0002 0.0005 0.001 0.002 0.003 0.006 0.01 0.02 0.04 0.07 0.1 0.15 0.2 0.25 0.3 0.4 0.5 0.6 0.7 0.8 0.9 1.0). For each individual λ state, the equilibration was performed for 1 ns ensemble, followed by 20 ns production run. The other parameters are the same as above.

## Acknowledgment

We thank Dr. Xuezhi Zhao and Dr. Dario Oliveira Passos for their helpful discussions. This study was supported by National Institutes of Health grants 5U54AI150472, and U01 AI136680 to DL and RL, P30 CA01495, R01 AI146017, and the Hearst Foundations to DL, and by a Bridge fund from Pace University to ND. AB was supported by the Margaret T. Morris Foundation and a fellowship from the Schmidt AI Futures. The calculations were run on the XSEDE allocation resource TGMCB100145 and a shared computing cluster at Temple University supported by the National Institutes of Health S10 OD020095.

## References

1. Engelman, A. N., Multifaceted HIV integrase functionalities and therapeutic strategies for their inhibition. J Biol Chem 2019, 294 (41), 15137–15157.

2. Di Santo, R., Inhibiting the HIV integration process: past, present, and the future. J Med Chem 2014, 57 (3), 539–66.

3. Puertas, M. C.; Ploumidis, G.; Ploumidis, M.; Fumero, E.; Clotet, B.; Walworth, C. M.; Petropoulos, C. J.; Martinez-Picado, J., Pan-resistant HIV-1 emergence in the era of integrase strand-transfer inhibitors: a case report. Lancet Microbe 2020, 1 (3), e130–e135.

4. Zhao, X. Z.; Smith, S. J.; Maskell, D. P.; Metifiot, M.; Pye, V. E.; Fesen, K.; Marchand, C.; Pommier, Y.; Cherepanov, P.; Hughes, S. H.; Burke, T. R., Jr., HIV-1 Integrase Strand Transfer Inhibitors with Reduced Susceptibility to Drug Resistant Mutant Integrases. ACS Chem Biol 2016, 11 (4), 1074–81.

5. Zhao, X. Z.; Smith, S. J.; Metifiot, M.; Marchand, C.; Boyer, P. L.; Pommier, Y.; Hughes, S. H.; Burke, T. R., Jr., 4-amino-1-hydroxy-2-oxo-1,8-naphthyridine-containing compounds having high potency against raltegravir-resistant integrase mutants of HIV-1. J Med Chem 2014, 57 (12), 5190–202.

6. Hare, S.; Gupta, S. S.; Valkov, E.; Engelman, A.; Cherepanov, P., Retroviral intasome assembly and inhibition of DNA strand transfer. Nature 2010, 464 (7286), 232–6.

7. Hare, S.; Smith, S. J.; Metifiot, M.; Jaxa-Chamiec, A.; Pommier, Y.; Hughes, S. H.; Cherepanov, P., Structural and functional analyses of the second-generation integrase strand transfer inhibitor dolutegravir (S/GSK1349572). Mol Pharmacol 2011, 80 (4), 565–72.

8. Metifiot, M.; Maddali, K.; Johnson, B. C.; Hare, S.; Smith, S. J.; Zhao, X. Z.; Marchand, C.; Burke, T. R., Jr.; Hughes, S. H.; Cherepanov, P.; Pommier, Y., Activities, crystal structures, and molecular dynamics of dihydro-1H-isoindole derivatives, inhibitors of HIV-1 integrase. ACS Chem Biol 2013, 8 (1), 209–17.

9. Hare, S.; Vos, A. M.; Clayton, R. F.; Thuring, J. W.; Cummings, M. D.; Cherepanov, P., Molecular mechanisms of retroviral integrase inhibition and the evolution of viral resistance. Proc Natl Acad Sci U S A 2010, 107 (46), 20057–62.

10. Smith, S. J.; Zhao, X. Z.; Passos, D. O.; Lyumkis, D.; Burke, T. R., Jr.; Hughes, S. H., Integrase Strand Transfer Inhibitors Are Effective Anti-HIV Drugs. Viruses 2021, 13 (2).

11. Passos, D. O.; Li, M.; Jóźwik, I. K.; Zhao, X. Z.; Santos-Martins, D.; Yang, R.; Smith, S. J.; Jeon, Y.; Forli, S.; Hughes, S. H.; Burke, T. R., Jr.; Craigie, R.; Lyumkis, D., Structural basis for strand-transfer inhibitor binding to HIV intasomes. Science (New York, N.Y.) 2020, 367 (6479), 810–814.

12. Sakae, Y.; Zhang, B. W.; Levy, R. M.; Deng, N., Absolute Protein Binding Free Energy Simulations for Ligands with Multiple Poses, a Thermodynamic Path That Avoids Exhaustive Enumeration of the Poses. J Comput Chem 2020, 41 (1), 56–68.

13. Raniolo, S.; Limongelli, V., Ligand binding free-energy calculations with funnel metadynamics. Nat Protoc 2020, 15 (9), 2837–2866.

14. Fu, H.; Shao, X.; Cai, W.; Chipot, C., Taming Rugged Free Energy Landscapes Using an Average Force. Acc Chem Res 2019, 52 (11), 3254–3264.

15. Khuttan, S.; Azimi, S.; Wu, J. Z.; Dick, S.; Wu, C.; Xu, H.; Gallicchio, E., Taming Multiple Binding Poses in Alchemical Binding Free Energy Prediction: the β-cyclodextrin Host-Guest SAMPL9 Blinded Challenge. ArXiv preprint 2023.

16. Arasteh, S.; Zhang, B. W.; Levy, R. M., Protein Loop Conformational Free Energy Changes via an Alchemical Path without Reaction Coordinates. J Phys Chem Lett 2021, 12 (18), 4368–4377.

17. Geronimo, I.; De Vivo, M., Alchemical Free-Energy Calculations of Watson-Crick and Hoogsteen Base Pairing Interconversion in DNA. J Chem Theory Comput 2022, 18 (11), 6966–6973.

18. Torrie, G. M.; Valleau, J. P., Nonphysical sampling distributions in Monte Carlo free-energy estimation: Umbrella sampling. Journal of Computational Physics 1977, 23 (2), 187–199.

19. Wang, F.; Landau, D. P., Efficient, Multiple-Range Random Walk Algorithm to Calculate the Density of States. Physical Review Letters 2001, 86 (10), 2050–2053.

20. Sun, Q.; Levy, R. M.; Kirby, K. A.; Wang, Z.; Sarafianos, S. G.; Deng, N., Molecular Dynamics Free Energy Simulations Reveal the Mechanism for the Antiviral Resistance of the M66I HIV-1 Capsid Mutation. Viruses 2021, 13 (5).

21. Gallicchio, E.; Levy, R. M., Recent theoretical and computational advances for modeling protein–ligand binding affinities. In Advances in Protein Chemistry and Structural Biology, Elsevier: 2011; Vol. 85, pp 27–80.

22. Deng, N.; Forli, S.; He, P.; Perryman, A.; Wickstrom, L.; Vijayan, S. K. V.; Tiefenbrunn, T.; Stout, C. D.; Gallicchio, E.; Olson, A. J.; Levy, R. M., Distinguishing Binders from False Positives by Free Energy Calculations: Fragment Screening Against the Flap Site of HIV Protease. Journal of Physical Chemistry B 2014, 119 (3), 976–988.

23. Wickstrom, L.; Gallicchio, E.; Chen, L.; Kurtzman, T.; Deng, N., Developing end-point methods for absolute binding free energy calculation using the Boltzmann-quasiharmonic model. Phys Chem Chem Phys 2022, 24 (10), 6037–6052.

24. Gerton, J. L.; Ohgi, S.; Olsen, M.; DeRisi, J.; Brown, P. O., Effects of mutations in residues near the active site of human immunodeficiency virus type 1 integrase on specific enzyme-substrate interactions. J Virol 1998, 72 (6), 5046–55.

25. Engelman, A.; Craigie, R., Identification of conserved amino acid residues critical for human immunodeficiency virus type 1 integrase function in vitro. J Virol 1992, 66 (11), 6361–9.

26. Jozwik, I. K.; Passos, D. O.; Lyumkis, D., Structural Biology of HIV Integrase Strand Transfer Inhibitors. Trends Pharmacol Sci 2020, 41 (9), 611–626.

27. Rhee, S. Y.; Grant, P. M.; Tzou, P. L.; Barrow, G.; Harrigan, P. R.; Ioannidis, J. P. A.; Shafer, R. W., A systematic review of the genetic mechanisms of dolutegravir resistance. J Antimicrob Chemother 2019, 74 (11), 3135–3149.

28. McColl, D. J.; Chen, X., Strand transfer inhibitors of HIV-1 integrase: bringing IN a new era of antiretroviral therapy. Antiviral Res 2010, 85 (1), 101–18.

29. Underwood, M. R.; Johns, B. A.; Sato, A.; Martin, J. N.; Deeks, S. G.; Fujiwara, T., The activity of the integrase inhibitor dolutegravir against HIV-1 variants isolated from raltegravir-treated adults. J Acquir Immune Defic Syndr 2012, 61 (3), 297–301.

30. Santoro, M. M.; Fornabaio, C.; Malena, M.; Galli, L.; Poli, A.; Menozzi, M.; Zazzi, M.; White, K. L.; Castagna, A.; Group, P. S., Susceptibility to HIV-1 integrase strand transfer inhibitors (INSTIs) in highly treatment-experienced patients who failed an INSTI-based regimen. Int J Antimicrob Agents 2020, 56 (1), 106027.

31. Cui, D.; Zhang, B. W.; Matubayasi, N.; Levy, R. M., The Role of Interfacial Water in Protein–Ligand Binding: Insights from the Indirect Solvent Mediated Potential of Mean Force. Journal of Chemical Theory and Computation 2018.

32. Zhang, B. W.; Cui, D.; Matubayasi, N.; Levy, R. M., The Excess Chemical Potential of Water at the Interface with a Protein from End Point Simulations. J Phys Chem B 2018, 122 (17), 4700–4707.

33. Cook, N. J.; Li, W.; Berta, D.; Badaoui, M.; Ballandras-Colas, A.; Nans, A.; Kotecha, A.; Rosta, E.; Engelman, A. N.; Cherepanov, P., Structural basis of second-generation HIV integrase inhibitor action and viral resistance. Science 2020, 367 (6479), 806–810.

34. Passos, D. O.; Li, M.; Jozwik, I. K.; Zhao, X. Z.; Santos-Martins, D.; Yang, R.; Smith, S. J.; Jeon, Y.; Forli, S.; Hughes, S. H.; Burke, T. R., Jr.; Craigie, R.; Lyumkis, D., Structural basis for strand-transfer inhibitor binding to HIV intasomes. Science 2020, 367 (6479), 810–814.

35. Summa, V.; Petrocchi, A.; Bonelli, F.; Crescenzi, B.; Donghi, M.; Ferrara, M.; Fiore, F.; Gardelli, C.; Gonzalez Paz, O.; Hazuda, D. J.; Jones, P.; Kinzel, O.; Laufer, R.; Monteagudo, E.; Muraglia, E.; Nizi, E.; Orvieto, F.; Pace, P.; Pescatore, G.; Scarpelli, R.; Stillmock, K.; Witmer, M. V.; Rowley, M., Discovery of raltegravir, a potent, selective orally bioavailable HIV-integrase inhibitor for the treatment of HIV-AIDS infection. J Med Chem 2008, 51 (18), 5843–55.

36. Krishnan, L.; Li, X.; Naraharisetty, H. L.; Hare, S.; Cherepanov, P.; Engelman, A., Structure-based modeling of the functional HIV-1 intasome and its inhibition. Proc Natl Acad Sci U S A 2010, 107 (36), 15910–5.

37. Zhao, X. Z.; Smith, S. J.; Maskell, D. P.; Metifiot, M.; Pye, V. E.; Fesen, K.; Marchand, C.; Pommier, Y.; Cherepanov, P.; Hughes, S. H.; Burke, T. R., Jr., Structure-Guided Optimization of HIV Integrase Strand Transfer Inhibitors. J Med Chem 2017, 60 (17), 7315–7332.

38. Li, M.; Oliveira Passos, D.; Shan, Z.; Smith, S. J.; Sun, Q.; Biswas, A.; Choudhuri, I.; Strutzenberg, T. S.; Haldane, A.; Deng, N.; Li, Z.; Zhao, X. Z.; Briganti, L.; Kvaratskhelia, M.; Burke, T. R., Jr.; Levy, R. M.; Hughes, S. H.; Craigie, R.; Lyumkis, D., Mechanisms of HIV-1 integrase resistance to dolutegravir and potent inhibition of drug-resistant variants. Sci Adv 2023, 9 (29), eadg5953.

39. Pronk, S.; Pall, S.; Schulz, R.; Larsson, P.; Bjelkmar, P.; Apostolov, R.; Shirts, M. R.; Smith, J. C.; Kasson, P. M.; van der Spoel, D.; Hess, B.; Lindahl, E., GROMACS 4.5: a high-throughput and highly parallel open source molecular simulation toolkit. Bioinformatics 2013, 29 (7), 845–854.

40. Tian, C.; Kasavajhala, K.; Belfon, K. A. A.; Raguette, L.; Huang, H.; Migues, A. N.; Bickel, J.; Wang, Y.; Pincay, J.; Wu, Q.; Simmerling, C., ff19SB: Amino-Acid-Specific Protein Backbone Parameters Trained against Quantum Mechanics Energy Surfaces in Solution. J Chem Theory Comput 2020, 16 (1), 528–552.

41. Ivani, I.; Dans, P. D.; Noy, A.; Pérez, A.; Faustino, I.; Hospital, A.; Walther, J.; Andrio, P.; Goñi, R.; Balaceanu, A.; Portella, G.; Battistini, F.; Gelpí, J. L.; González, C.; Vendruscolo, M.; Laughton, C. A.; Harris, S. A.; Case, D. A.; Orozco, M., Parmbsc1: a refined force field for DNA simulations. Nature Methods 2015.

42. Wang, J. M.; Wolf, R. M.; Caldwell, J. W.; Kollman, P. A.; Case, D. A., Development and testing of a general amber force field. Journal of Computational Chemistry 2004, 25 (9), 1157–1174.

43. Jakalian, A.; Bush, B. L.; Jack, D. B.; Bayly, C. I., Fast, efficient generation of high-quality atomic Charges. AM1-BCC model: I. Method. Journal of Computational Chemistry 2000, 21 (2), 132–146.

44. Sun, S.; Huggins, D. J., Assessing the effect of forcefield parameter sets on the accuracy of relative binding free energy calculations. Front Mol Biosci 2022, 9, 972162.

45. Sun, Z.; Zhang, J. Z. H., Thermodynamic Insights of Base Flipping in TNA Duplex: Force Fields, Salt Concentrations, and Free-Energy Simulation Methods. CCS Chemistry 2021, 3 (2), 1026–1039.

46. Wickstrom, L.; Deng, N.; He, P.; Mentes, A.; Nguyen, C.; Gilson, M. K.; Kurtzman, T.; Gallicchio, E.; Levy, R. M., Parameterization of an effective potential for protein-ligand binding from host-guest affinity data: Force Field Optimization With Host-Guest Systems. Journal of Molecular Recognition 2016, 29 (1), 10–21.

47. Li, P.; Merz, K. M., Jr., Metal Ion Modeling Using Classical Mechanics. Chem Rev 2017, 117 (3), 1564–1686.

48. Li, P.; Merz, K. M., Jr., MCPB.py: A Python Based Metal Center Parameter Builder. J Chem Inf Model 2016, 56 (4), 599–604.

49. Vedani, A.; Huhta, D. W., A new force field for modeling metalloproteins. Journal of the American Chemical Society 1990, 112 (12), 4759–4767.

50. Lin, F.; Wang, R., Systematic Derivation of AMBER Force Field Parameters Applicable to Zinc-Containing Systems. J Chem Theory Comput 2010, 6 (6), 1852–70.

51. Seminario, J. M., Calculation of intramolecular force fields from second-derivative tensors. International Journal of Quantum Chemistry 1996, 60 (7), 1271–1277.

52. Melse, O.; Antes, I.; Kaila, V. R. I.; Zacharias, M., Benchmarking biomolecular force field-based Zn2+ for mono- and bimetallic ligand binding sites. J Comput Chem 2023, 44 (8), 912–926.

53. He, P.; Zhang, B. W.; Arasteh, S.; Wang, L.; Abel, R.; Levy, R. M., Conformational Free Energy Changes via an Alchemical Path without Reaction Coordinates. J Phys Chem Lett 2018, 9 (15), 4428–4435.

54. Berendsen, H. J. C. a. P., J. P. M. and van Gunsteren, W. F. and DiNola, A. and Haak, J. R., Molecular dynamics with coupling to an external bath The Journal of chemical physics 1984, 81, 3684–3690.

55. Essmann, U.; Perera, L.; Berkowitz, M. L.; Darden, T.; Lee, H.; Pedersen, L. G., A Smooth Particle Mesh Ewald Method. Journal of Chemical Physics 1995, 103 (19), 8577–8593.

