## Supporting Information for "Elucidating the molecular determinants for binding modes of a third-generation HIV-1 integrase strand transfer inhibitor: Importance of side chain and solvent reorganization"

11/25/2023

*Qinfang Sun[1], Avik Biswas[2,3], Dmitry Lyumkis[2,4], Ronald Levy[1], Nanjie Deng[5]**

(1) Center for Biophysics and Computational Biology and Department of Chemistry, Temple University, Philadelphia, PA 19122

(2) The Salk Institute for Biological Studies, Laboratory of Genetics, La Jolla, CA 92037

(3) Department of Physics, University of California San Diego, La Jolla, CA, 92093

(4) Graduate schools for Biological Sciences, Section of Molecular Biology, University of California, San Diego, La Jolla, CA, 92093

(5) Department of Chemistry and Physical Sciences, Pace University, New York, NY10038

Table S1. RMSD in the positions of C-alpha atoms between the corresponding binding site residues (within 5 Å from any 4f atoms) in the HIV-1 and PFV intasomes. For the two DNA nucleotides that form part of the binding site, the corresponding deviations between the backbone C5’ atoms are shown.

| Corresponding residue pairs | RMSD (Å) |
|---|---|
| N117(HIV)/Q186(PFV) | 0.39 |
| Y143(HIV)/Y212(PFV) | 0.36 |
| P145(HIV)/P214(PFV) | 0.38 |
| Q146(HIV)/Q215(PFV) | 0.42 |
| Q148(HIV)/S217(PFV) | 0.41 |
| DNA: A36(HIV)/A16(PFV) | 0.43 |
| DNA: C35(HIV)/C17(PFV) | 1.06 |

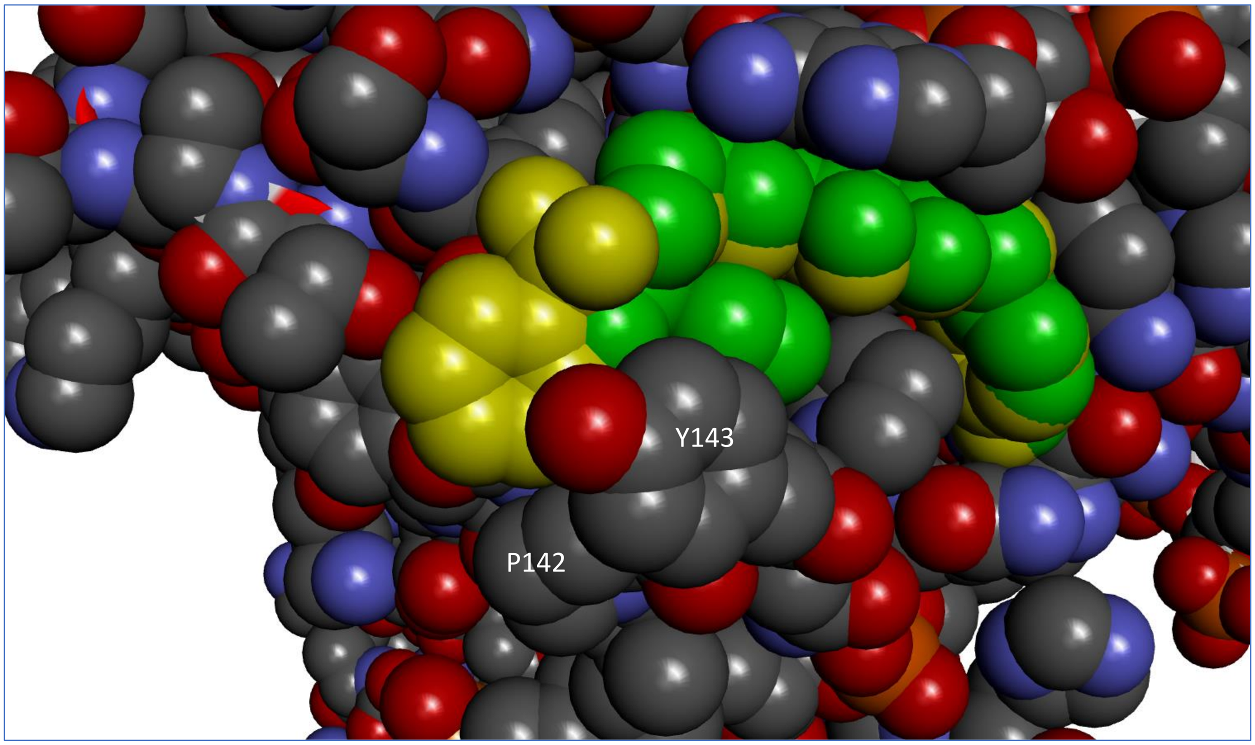


*Figure S1. Extended (yellow) and bent (green) 4f in the binding site of HIV-1 intasome. To facilitate a conformational transition between the two binding conformations requires both side chain atoms from the protein residues P142 and Y143 to rearrange to avoid steric clash with the sulfonylphenyl moiety of 4f converting from the extended to the bent conformation and vice versa.*

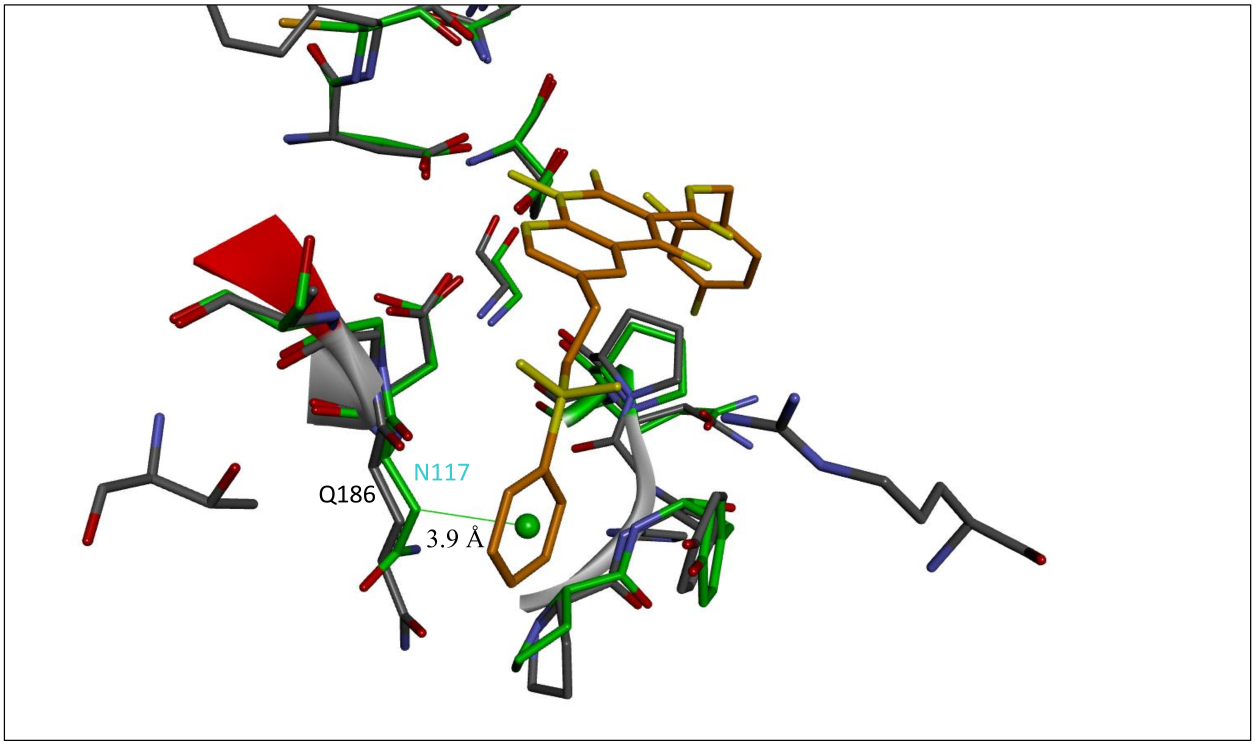


*Figure S2. Nonpolar interactions between the CB of N117 of HIV-1 and Q186 of PFV with the sulfonylphenyl moiety of the extended 4f. The green sphere represents the centroid of the terminal benzene ring of 4f.*

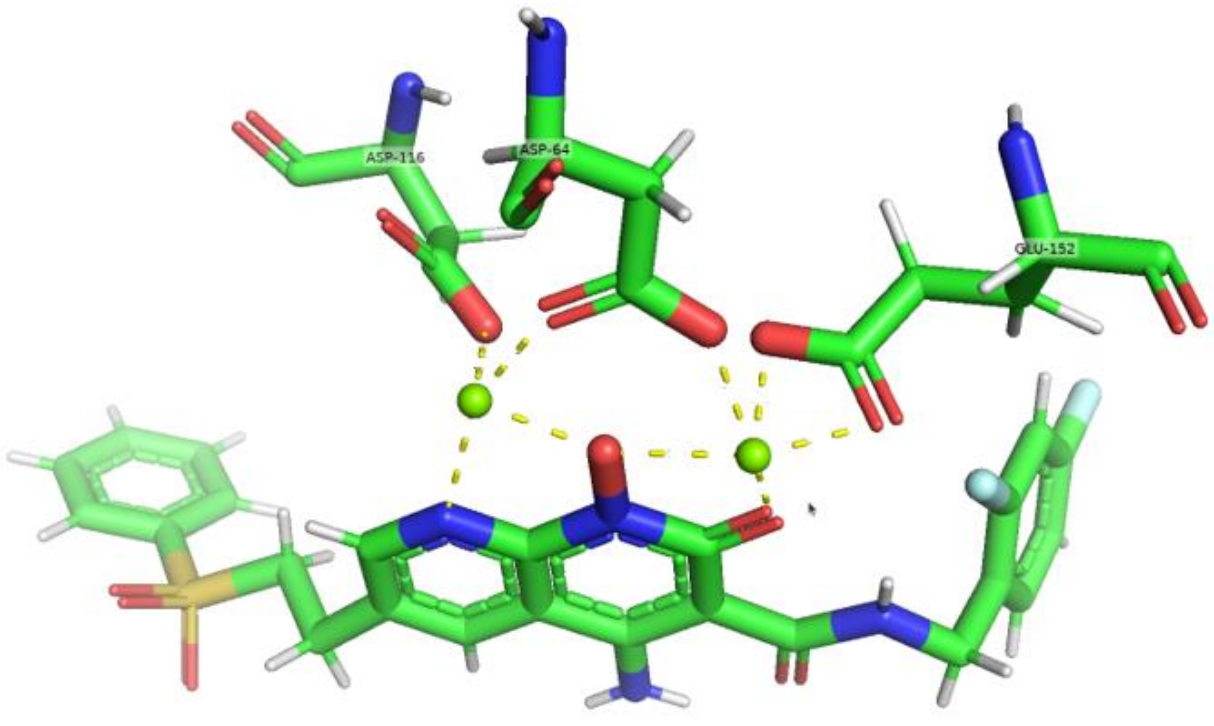


*Figure S3. The HIV-1 IN active site carboxylates Asp64, Asp116, and Glu152 coordinate a pair of $Mg^{2+}$ ions, which in turn interact with the metal-chelating core naphthyridine ring of the INSTIs 4f (bottom molecule)*

Table S2. The bond, angle, corresponding force constants, the partial charge, and the VDW parameters generated by MCPB.py program with GAMESS-US for $Mg^{2+}$ associated atoms in HIV-1 and PFV intasomes.

| Bond parameters for Different $Mg^{2+}$ | Atom1 Name | | Atom2 Name | R eq(Å) | Force Const. (kcal/mol*Å^2) |
|---|---|---|---|---|---|
| HIV-1 MG1 | OD1 | | MG | 1.9747 | 73.8 |
| | OD2 | | MG | 2.0337 | 56.7 |
| | MG | | OAF | 2.04 | 63.4 |
| | MG | | NAV | 2.2535 | 34 |
| HIV-1 MG2 | OD2 | | MG | 1.9886 | 66 |
| | OE2 | | MG | 2.0385 | 56 |
| | OE1 | | MG | 2.0739 | 45.5 |
| | MG | | OAC | 2.0893 | 47.9 |
| | MG | | OAF | 2.0794 | 48.7 |
| PFV MG1 | OD1 | | MG | 1.9746 | 71.5 |
| | OD2 | | MG | 2.0484 | 51.6 |
| | MG | | OAF | 2.0398 | 63 |
| | MG | | NAV | 2.2665 | 28.7 |
| PFV MG2 | OD2 | | MG | 1.9837 | 69.4 |
| | OE2 | | MG | 2.044 | 53.2 |
| | OE1 | | MG | 2.0703 | 46.5 |
| | MG | | OAC | 2.0886 | 46.5 |
| | MG | | OAF | 2.0785 | 48.9 |
| Angle parameters for Different $Mg^{2+}$ | Atom1 Name | Atom2 Name | Atom3 Name | Theta eq (Rad) | Force Const. (kcal/mol*Rad^2) |
| HIV-1 MG1 | OD1 | MG | OD2 | 115.51 | 32.65 |
| | OD1 | MG | OAF | 91.96 | 57.16 |
| | OD1 | MG | NAV | 132.3801 | 34.49 |
| | CG | OD1 | MG | 137.3601 | 51.76 |
| | OD2 | MG | OAF | 117.5801 | 56.9 |
| | OD2 | MG | NAV | 111.37 | 64.84 |
| | CG | OD2 | MG | 90.35 | 112.53 |
| | MG | OAF | NBI | 118.9801 | 73.23 |
| | MG | OAF | MG | 125.8301 | 54.8 |
| | MG | NAV | CAP | 120.7801 | 94.52 |
| | MG | NAV | CBH | 120.7801 | 94.52 |
| | OAF | MG | NAV | 73.27 | 53.5 |
| | OD2 | MG | OE1 | 69.57 | 103.97 |
| | OD2 | MG | OE2 | 31.76 | 116.8201 |

| | | | | | |
|---|---|---|---|---|---|
| HIV-1 MG2 | OD2 | MG | OAC | 36.62 | 133.0401 |
| | OD2 | MG | OAF | 59.84 | 89.03 |
| | CG | OD2 | MG | 55.29 | 141.3901 |
| | OE2 | MG | OAC | 52 | 110.08 |
| | OE2 | MG | OAF | 71.21 | 113.85 |
| | OE1 | MG | OE2 | 127.45 | 63.26 |
| | OE1 | MG | OAC | 91.26 | 94.42 |
| | OE1 | MG | OAF | 52.12 | 166.6201 |
| | CD | OE1 | MG | 125.83 | 87.84 |
| | CD | OE2 | MG | 119.79 | 89.29 |
| | MG | OAF | MG | 54.8 | 125.8301 |
| | MG | OAC | CBF | 55.89 | 118.7601 |
| | MG | OAF | NBI | 84.05 | 114.04 |
| | OAC | MG | OAF | 50.48 | 74 |
| PFV MG1 | OD1 | MG | OD2 | 116.34 | 32.47 |
| | OD1 | MG | OAF | 90.14 | 56.2 |
| | OD1 | MG | NAV | 143.9901 | 39.35 |
| | CG | OD1 | MG | 140.1601 | 52.54 |
| | OD2 | MG | OAF | 121.5801 | 47.21 |
| | OD2 | MG | NAV | 99.49 | 51.05 |
| | CG | OD2 | MG | 89.46 | 118.25 |
| | MG | OAF | NBI | 119.3901 | 60.94 |
| | MG | OAF | MG | 126.8701 | 49.9 |
| | MG | NAV | CAP | 120.6801 | 98.36 |
| | MG | NAV | CBH | 120.6801 | 98.36 |
| | OAF | MG | NAV | 72.93 | 57.55 |
| PFV MG2 | OD2 | MG | OE1 | 104.32 | 70.41 |
| | OD2 | MG | OE2 | 116.4 | 17.21 |
| | OD2 | MG | OAC | 137.5401 | 17.75 |
| | OD2 | MG | OAF | 89.38 | 63.27 |
| | CG | OD2 | MG | 139.0801 | 33.99 |
| | OE2 | MG | OAC | 106.06 | 43.29 |
| | OE2 | MG | OAF | 112.39 | 61.5 |
| | OE1 | MG | OE2 | 63.26 | 129.05 |
| | OE1 | MG | OAC | 94.19 | 84.98 |
| | OE1 | MG | OAF | 166.1901 | 62.78 |
| | CD | OE1 | MG | 88.03 | 125.33 |
| | CD | OE2 | MG | 89.06 | 119.33 |
| | MG | OAF | MG | 126.8701 | 49.9 |

| | | | | | |
|---|---|---|---|---|---|
| | MG | OAC | CBF | 118.4101 | 54.8 |
| | MG | OAF | NBI | 113.72 | 76.6 |
| | OAC | MG | OAF | 74.06 | 63.1 |
| VdW parameters and charge for $Mg^{2+}$ | LJ Radius/Å | LJ Depth/kcal/mol | Mass | Charge | |
| HIV-1 MG1 | 1.373 | 0.0118 | 24.3 | 1.5414 | |
| HIV-1 MG2 | 1.373 | 0.0118 | 24.3 | 1.6172 | |
| PFV MG1 | 1.373 | 0.0118 | 24.3 | 1.508 | |
| PFV MG2 | 1.373 | 0.0118 | 24.3 | 1.542 | |
